# Active elimination of intestinal cells drives oncogenic growth in organoids

**DOI:** 10.1101/2020.11.14.378588

**Authors:** Ana Krotenberg Garcia, Arianna Fumagalli, Huy Quang Le, Owen J. Sansom, Jacco van Rheenen, Saskia J.E. Suijkerbuijk

## Abstract

Competitive cell-interactions play a crucial role in quality control during development and homeostasis. Here we show that cancer cells use such interactions to actively eliminate wild-type intestine cells in enteroid monolayers and organoids. This apoptosis-dependent process boosts proliferation of intestinal cancer cells. The remaining wild-type population activates markers of primitive epithelia and transits to a fetal-like state. Prevention of this cell fate transition avoids elimination of wild-type cells and, importantly, limits the proliferation of cancer cells. JNK signalling is activated in competing cells and is required for cell fate change and elimination of wild-type cells. Thus, cell competition drives growth of cancer cells by active out-competition of wild-type cells through forced cell death and cell fate change in a JNK dependent manner.

## Introduction

Over the past years it became evident that the internal proliferative potential of tumor cells is not sufficient for their expansion. Instead, tumor cells need to acquire multiple hallmarks of cancer, including growth-supporting interplay of tumor cells and their environment in order to sustain their proliferation (Hanahan and Weinberg, 2011). The basis of this interaction is often formed by processes that are, in origin, essential for normal early development and homeostasis (Suijkerbuijk and van Rheenen, 2017). One of those processes, cell competition, regulates survival of cells based on their relative fitness. In a homotypic context, cells strive and form viable tissues. However, in tissues built by heterogenous populations weaker cells will be removed by surrounding stronger cells. These features provide a strong mechanism that controls overall tissue and organismal fitness (Bowling et al., 2019; Clavería and Torres, 2016). Indeed, quality control by cell competition starts in the early mouse embryo (Clavería et al., 2013; Sancho et al., 2013) and continuous to impact physiology up to late adulthood by determining the speed of aging (Merino et al., 2015).

In a tumor context, it has been shown that relative activation of YAP/TAZ in peritumoral hepatocytes can influence growth of liver tumors by a process akin to cell competition (Moya et al., 2019). Furthermore, entosis, a form of cancer-driven cell competition, is correlated with a poor prognosis in patients with pancreatic ductal adenocarcinoma (Hayashi et al., 2020). These are examples of passive types of competitive cell interactions that suggest that cell competition could influence oncogenic growth. In addition, active cell competition enforced by differential expression of isoforms of the protein Flower, gives human cancer cells a competitive advantage over surrounding stromal tissue (Madan et al., 2019). We have shown that adenomas in the *Drosophila* midgut are dependent on active elimination of healthy surrounding tissue for their colonization (Suijkerbuijk et al., 2016). This illustrates that oncogenic growth can be driven by active cell competition. However, the full potential and most of the mechanisms behind this process still need to be uncovered.

Here, we report that cell competition promotes growth of cancer cells in intestinal organoids. We show that cancer cells utilize JNK signaling to actively eradicate wild-type intestinal cells. Upon competition, JNK activation promotes a cell fate transition in wild-type cells, which revert to a fetal-like state that is normally observed upon acute injury (Gregorieff et al., 2015; Nusse et al., 2018; Yui et al., 2018). Together, these competitional processes result in an increased colonization potential of cancer cells.

## Results

### Wild-type small intestine cells are eliminated by cancer cells in enteroid monolayers

In order to investigate whether cell competition plays a role in mammalian intestinal cancer, we adapted the recently described enteroid monolayer culture system that recapitulates all key aspects of the intestinal epithelium (Thorne et al., 2018). Two different types of organoid cultures were derived from mouse small intestines: membrane-bound tdTomato-labeled wild-type cells and Dendra2-labeled intestinal cancer cells, derived from Villin-Cre^ERT2^:*Apc*^fl/fl^*Kras*^G12D/WT^*Trp53*^fl/R172H^ transgenic mice (Fumagalli et al., 2017a). Next, dissociated single cells from these cultures were plated separately or mixed together on matrix coated imaging plates and followed for up to 10 days (Figure 1A). Pure cultures gradually covered the surface until a stable enteroid monolayer was formed (Figures 1B-1D). However, once a full monolayer developed in mixed culture conditions, the surface area taken up by wild-type cells was not maintained but instead gradually decreased over time (Figures 1B’ and 1C). This implies that cancer cells can out-compete wild-type cells in mixed enteroid monolayer cultures. To further study the fate of wild-type cells in mixed cultures we used time-lapse imaging. Interestingly, wild-type cells decreased both in size and number when mixed with cancer cells (Figure 1E; Videos S1 and S2), suggesting that wild-type cells are compressed and eventually eliminated by tumor cells. Consistently, we observed higher rates of cleaved-Caspase 3, a marker of apoptosis, in competing wild-type cells (Figure 1F). Together, these findings indicate that fitter cancer cells take over enteroid monolayers at the expense of less fit wild-type intestinal cells, which is a hallmark of cell competition (Morata and Ripoll, 1975).

**Figure 1.**
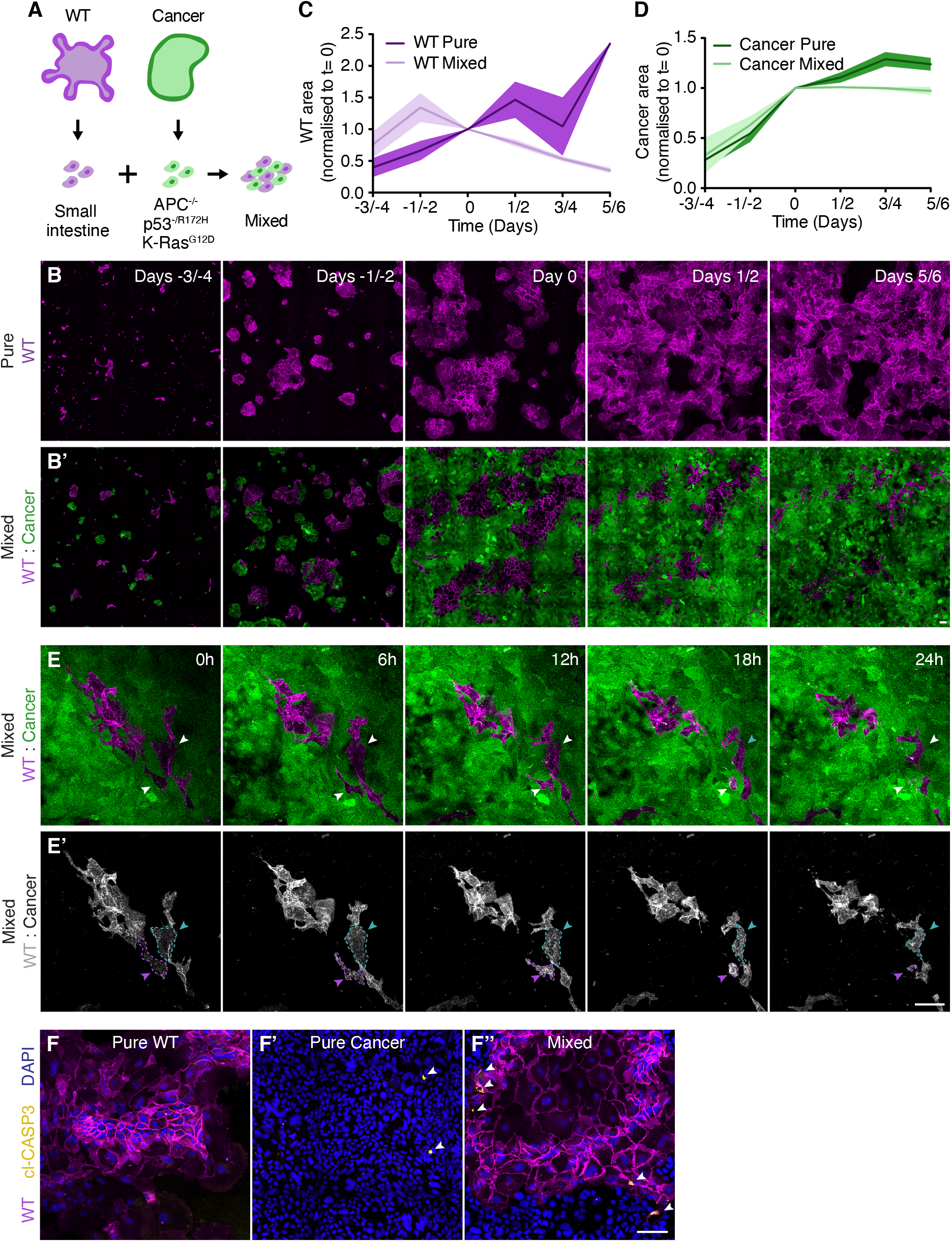
Wild-type small intestine cells are eliminated by cancer cells in enteroid monolayers. A) Schematic representation of a model for cell competition in murine enteroid monolayers. B-D) Representative pictures of sequential imaging of enteroid monolayers in pure (B) and mixed (B’) conditions and quantification of the surface covered by wild-type (C) or cancer (D) populations over time normalized to Day 0 (Mean ±SEM). Day 0 is the moment a full monolayer is formed in mixed conditions. E) Representative images of time-lapse series of a competing enteroid monolayer, arrow heads indicate examples of wild-type cells that are shrinking (cyan) and being eliminated (magenta). F) Representative confocal images of pure (F and F’) and mixed (F’’) enteroid monolayers. Apoptotic cells are marked by cleaved-caspase 3 (yellow) and indicated by arrow heads. Scale bars = 100μm.

### Cancer cells outcompete wild-type small intestine cells

In order to study the interaction between cancer cells and wild-type cells in near-native conditions, we exploited the 3D organoid system (Sato et al., 2009), which closely resembles the architecture of the intestinal tissue. Wild-type and cancer cell cultures were dissociated into small clumps of cells and concentrated to enable formation of mixed organoids (Figure 2A). Time-lapse imaging of these cultures revealed that whereas pure wild-type cultures could expand over time (Figure 2B; Video S3), wild-type cells in mixed structures gradually disappeared (Figure 2B’; Video S4). Interestingly, two morphological changes occurred to wild-type cells in mixed organoids compared to in pure organoids, the formation of typical crypt-villus structures was severely diminished and the extrusion of wild-type cells into the lumen of organoids increased (Figure 2B). Together, these data suggest that the presence of cancer cells within the organoid directly affects the behavior of wild-type cells. Next, we followed whole drops of matrix containing wild-type or mixed organoids for up to three days. Quantification of the surface occupied by different populations in organoids showed that wild-type cells present in mixed organoids did not increase over time, whilst wild-type cells in pure organoids or the cancer cell population in both types of organoids rapidly expanded (Figures 2C-2E). Thus, similar to enteroid monolayers, wild-type cells are outcompeted by cancer cells in near-native 3D organoid structures.

**Figure 2.**
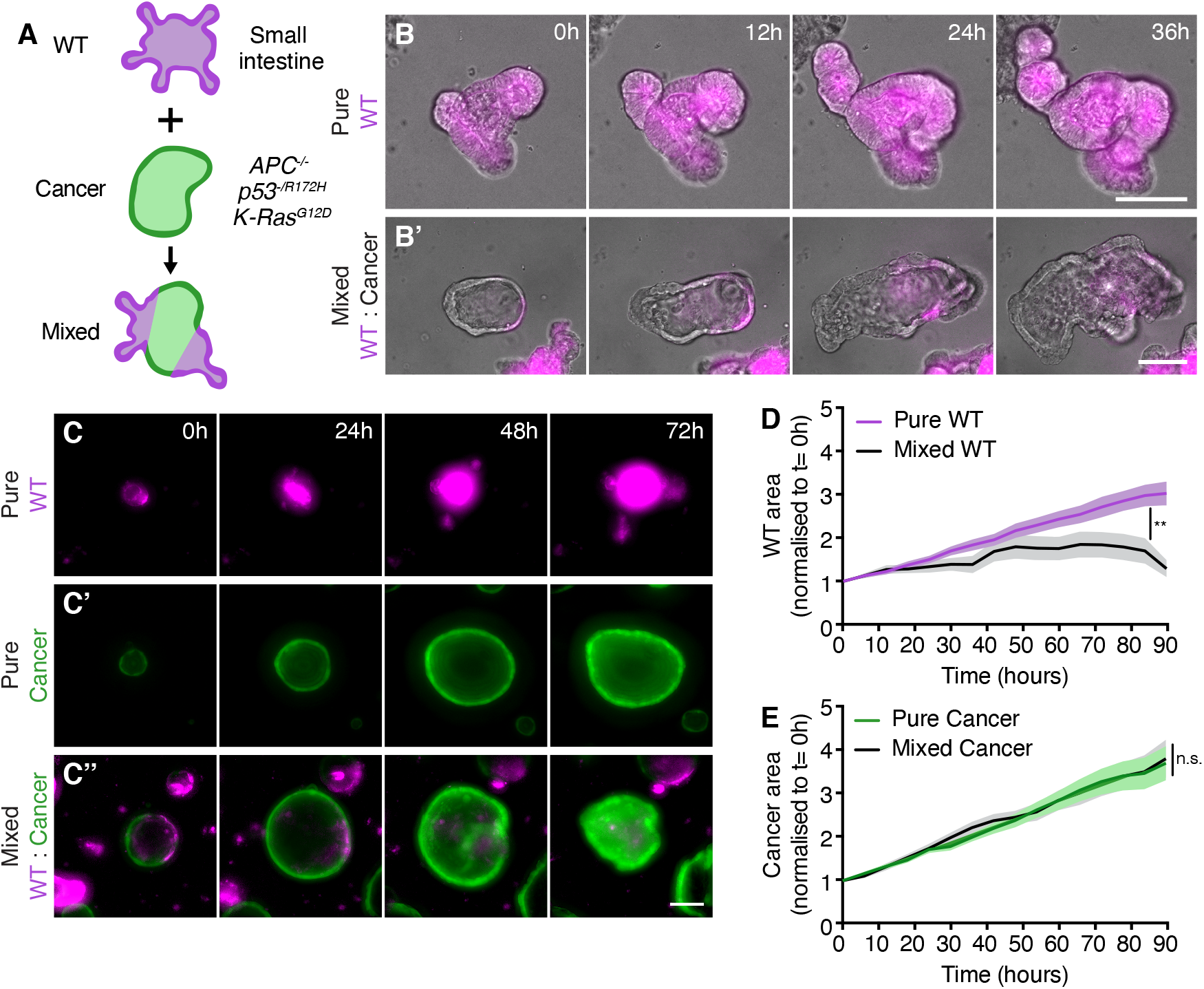
Cancer cells outcompete wild-type small intestine cells. A) Schematic depiction of a 3D model for cell competition in murine intestinal organoids. B) Representative images of time-lapse series of pure wild-type (B) and mixed (B’) intestinal organoids. C-E) Analysis of wild-type and cancer organoid growth under pure and mixed conditions by live-imaging. Representative images (C) and quantification of the area covered by wild-type (D) or cancer (E) cells within organoids normalized to the start of the time-lapse (Mean ±SEM, Unpaired t-test, two-tailed; p= 0.0074, n= 22 & 22 organoids (D); p=0.8897, n= 30 & 30 organoids (E)). Scale bars = 100μm.

### Direct interaction is essential for cell competition

Using higher resolution imaging we next investigated the cellular interactions formed by different cell populations in organoids. Actin filaments of the cytoskeleton, visualized using Phalloidin, revealed close cell-cell interactions within pure organoids (Figures 3A and 3A’). Importantly, these close cell-cell contacts were similar in mixed organoids (Figures 3A’’ and 3A’’’; Video S5). Furthermore, wild-type cells and cancer cells intermingled and shared cell contacts, showing that competing cells are part of the same epithelium and that these competing cell populations directly interact.

**Figure 3.**
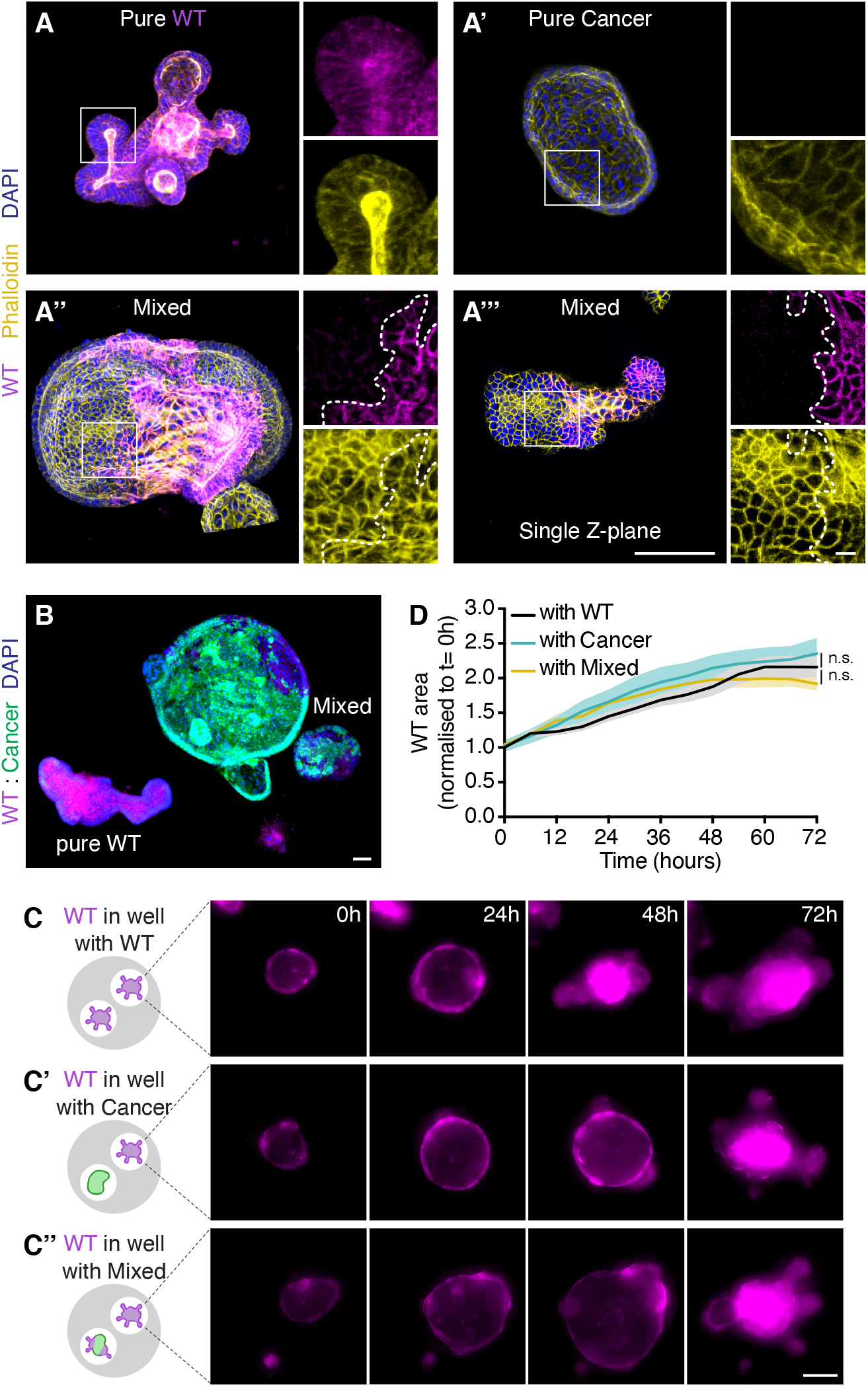
Direct interaction is essential for cell competition. A) Representative 3D-reconstructed confocal images of pure WT (A), pure cancer (A’) and mixed (A’’) organoids, and a single Z-plane of A’’ (A’’’). The actin cytoskeleton is stained with Phalloidin (yellow), nuclei with DAPI (blue) and borders between wild-type and cancer cells are indicated by dashed lines. The insets display a 2.5x magnification of the area in the white box. B) Representative confocal image of a mixed culture containing a pure WT organoid, nuclei are visualized with DAPI (blue). C-D) Representative images from live-imaging of pure WT organoids co-cultured with pure WT (C), pure cancer (C’) or mixed (C’’) organoids and quantification of the area covered by wild-type cells within indicated organoids (D) normalized to the start of the time-lapse (Mean ±SEM, 2-way ANOVA, multiple comparisons, n=18 organoids for each condition, ‘WT in WT’ vs. ‘WT in cancer’ p= 0.5453, ‘WT in WT’ vs. ‘WT in Mix’ p= 0.9689). Scale bars = 100μm, excluding magnifications in (A) where scale bar = 10μm.

We observed that occasional organoids of pure wild-type origin could persist in drops of matrix that were occupied by mixed structures (Figure 3B). Suggesting that elimination of wild-type cells depends on their direct interaction with cancer cells. In order to further characterize this observation, we plated pure wild-type organoids in the same well together with pure wild-type, pure cancer or mixed organoids and followed these drops for multiple days. Neither the presence of pure cancer nor mixed cultures affected the growth or phenotype of wild-type organoids within the same well (Figures 3C and 3D). Thus, cell competition is short-ranged and the presence of both cell populations within the same organoid is required for out-competition of wild-type cells. In addition, it is important to note that this shows that neither factors secreted by pure cancer cells or mixed organoids nor depletion of components in the growth medium are sufficient to induce elimination of wild-type cells. Together, these data suggest that cell competition driven by cancer cells is active and short-ranged.

### Elimination of wild-type cells is driven by apoptosis

So far, we observed that the surface area of wild-type cells that contribute to mixed organoids declines over time. This is supported by a reduction in the percentage of wild-type cells in organoids from ±40% one day after mixing to ±10% on day four (Figures 4A and 4B). Additionally, we observed that the absolute number of wild-type cells that contribute to mixed organoids was lower four days after mixing while the number of total cells and of cancer cells increased (Figure 4C). This indicates that, although expansion of the wild-type cell population is slower than that of the cancer cell population, a difference in proliferation rate cannot be the sole determinant of the loss of wild-type cells.

**Figure 4.**
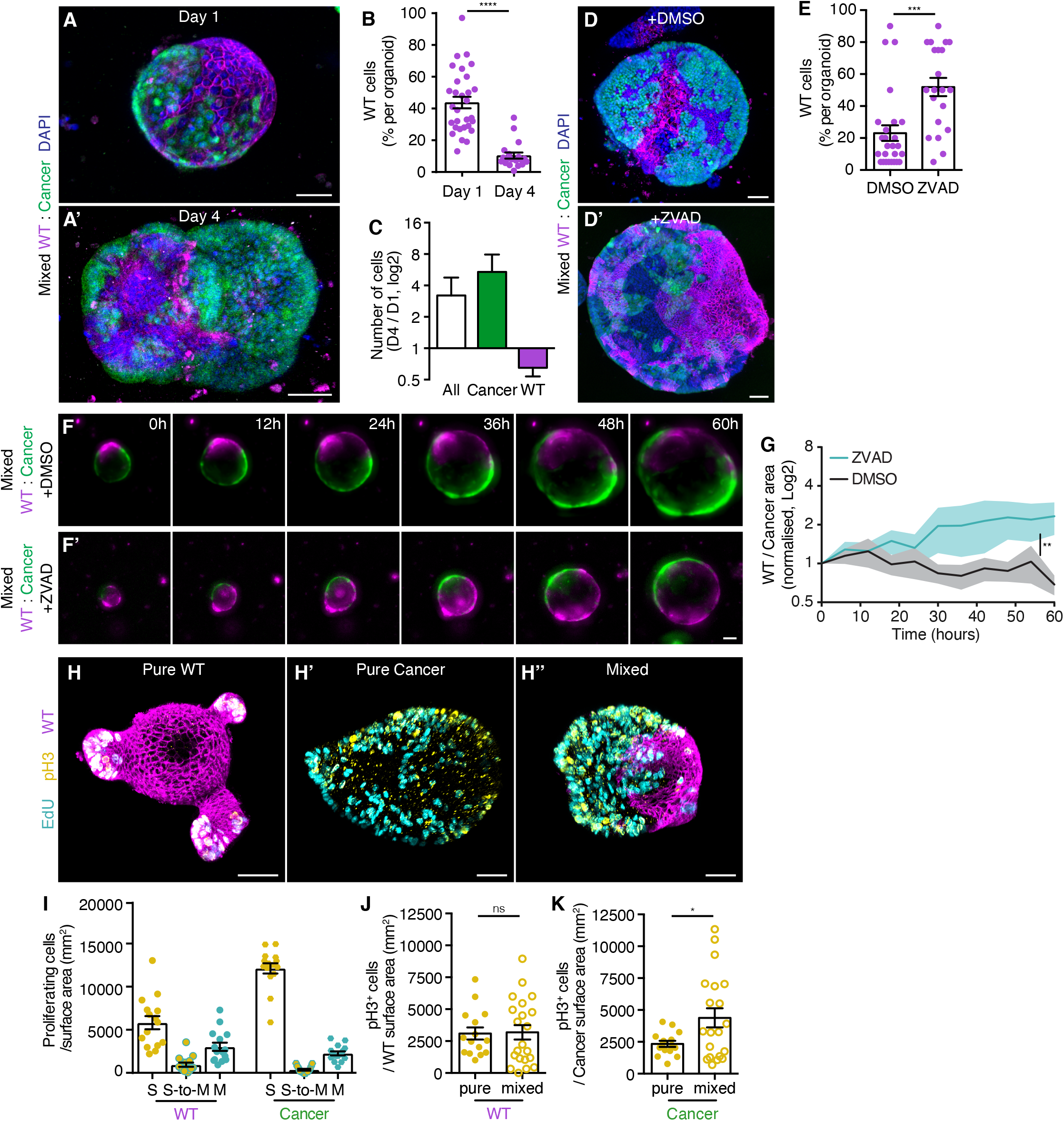
Elimination of wild-type cells is driven by apoptosis. A) Representative 3D-reconstructed confocal images of mixed organoids 1 day (A) and 4 days (A’) after plating, nuclei are stained with DAPI (blue). B) The percentage of wild-type cells contributing to mixed organoids on Day 1 and Day 4 after plating is shown, each dot represents one organoid (Mean ±SEM, Mann-Withney, two-tailed, p<0.0001, n= 30 &19 organoids). C) Displays the absolute number of cells in organoids shown in ‘B’, the ratio of ‘day 4’ over ‘day 1’ of all (white), cancer (green) and wild-type (magenta) cells are plotted on a Log2 scale (Mean ±SEM). D-E) Representative 3D-reconstructed confocal images of control (D) and apoptosis inhibited (D’) mixed organoids 4 days after plating, nuclei are stained with DAPI (blue), and quantification of the percentage of wild-type cells contributing to mixed organoids 4 days after plating (E), each dot represents one organoid (Mean ±SEM, Mann-Withney, two-tailed, p=0.0004, n= 26 &21 organoids). F-G) Representative images of control (F) and Z-VAD-FMK (F’) treated mixed organoids by live-imaging and quantification of the ratio of the area of wild-type / cancer cells (G) in mixed organoids normalized to the start of the time-lapse on a Log2 scale (Mean ±SEM, 2-way Anova Multiple Comparisons, two-tailed, p=0.0001, n= 24 &24). H-I) Representative 3D-reconstructed confocal images of pure wild-type (H), pure cancer (H’) and mixed organoids 3 days after plating and quantification of the number of cells in the different stages of the cell cycle per pure wild-type (left bars) and cancer (right bars) surface area (I). Cells in S-phase are labelled with EdU (cyan) and mitotic cells are marked by pH3 (yellow), cells that progressed from S-phase to mitosis within one hour are double positive. Each dot in (I) represents one organoid (Mean ±SEM, n= 15 organoids for each condition). J-K) Graphs display the number of mitotic cells per wild-type (J) and cancer (K) surface area in pure and mixed conditions, each dot represents one organoid (Mean ±SEM, unpaired T-test, two-tailed; p=0.9041, n=15 & 21 organoids (J); p=0.0288, n=15 & 20 organoids (K)). Scale bars = 50μm

A major hallmark of cell competition is the induction of cell death in weaker cells (Amoyel and Bach, 2014). In order to characterize how wild-type cells are removed from mixed organoids, we next asked if programmed cell death is involved. Treatment with the pan-Caspase inhibitor Z-VAD-FMK resulted in maintenance of a substantial population of wild-type cells in mixed organoids four days after mixing (Figures 4D and 4E). Furthermore, time-lapse imaging of Z-VAD-FMK treated cultures confirmed that wild-type cells were not eliminated (Figures 4F and 4G), while expansion of pure cultures was not affected (Figures S1A and S1B). Interestingly, the relative contribution of wild-type cells, but not the total surface area of mixed organoids was altered upon treatment with Z-VAD-FMK (Figure S1C). Thus, together these data show that out-competition of wild-type cells is active and dependent on programmed cell death.

### Cancer cells boost their growth by cell competition

Next, we questioned whether cancer cells could in turn be affected by the presence of wild-type cells. First, we determined the basal proliferation rates of both wild-type and cancer cells in unchallenged conditions. Using markers that identify DNA replication (1-hour EdU pulse) and active cell division (pH3), we could distinguish three populations of cells; cells in S-phase (S, EdU+), cells that proceeded from S-phase to mitosis (S-to-M, EdU+/pH3+) and mitotic cells (M, pH3+) (Figure 4F). Whereas proliferation in pure wild-type organoids was restricted to crypt regions, no obvious spatial organization of proliferating cells was observed in pure cancer organoids (Figures 4F’ and 4F’’). Furthermore, cells followed the expected distribution over different stages of the cell cycle in relation of the duration of these stages (S>M>S-to-M; Figure 4I). It is important to note that although the number of cells in S-phase is higher among cancer cells than among wild-type cells, the percentage of those cells that proceeded to mitosis within one hour was significantly lower in cancer (4%) compared to wild-type (18%) cells (p= 0.0007). This suggests that the EdU incorporation is not necessarily a reflection of proliferation rates, and is also affected by S-phase length in these cultures. Therefore, we decided to use the mitotic index, based on pH3 staining as a proxy for cell proliferation. No altered cell division rate was observed in wild-type cells three days after mixing with cancer cells, indicating that those cells that are not eliminated can still proliferate (Figure 4J). In contrast, cancer cells showed a marked increased mitotic index in competing conditions (Figure 4K). This implies that intestinal cancer cells boost their own proliferation and benefit from competitive cell interactions with wild-type intestinal cells.

#### Cell competition induces a fetal-like state in WT cells

In order to characterize molecular mechanisms underlying cell competition driven by cancer cells we used bulk RNA sequencing to identify genes that are differentially expressed between pure and mixed organoids (Figures S2A-S2C). The transcriptome of competing cancer cells was very similar to that of pure cancer cells (Figure S2D), indicating that phenotypic changes in cancer cells induced by cell competition are not of a transcriptional nature. In contrast, the transcriptome of wild-type cells was dramatically changed upon cell competition (Figure S2D). Subsequent Gene Ontology analysis displayed enrichment of multiple cell death related pathways in competing wild-type cultures (Figure 5A) whereas processing of mRNA and cell proliferation were enriched in pure wild-type cultures (Figure 5B). These data confirm a negative impact of cell competition on wild-type cells. Interestingly, among the most highly upregulated genes were multiple members of the *Ly6* family (Figure 5C). The *Ly6* family genes are induced in intestinal epithelia after exposure to colitis (Flanagan et al., 2008) and one of its members, Stem Cell Antigen-1 (*Sca1/Ly6a*), has recently been shown to be a marker of regenerating colonic epithelia (Yui et al., 2018). Furthermore, *Sca1* expression is activated in small intestinal epithelia that have been challenged with parasitic helminths (Nusse et al., 2018). Importantly, both injury responses cause a reprogramming of the tissue and adoption of a fetal-like state (Nusse et al., 2018; Yui et al., 2018). This response is also characterized by increased expression of genes of the Annexin family (Yui et al., 2018), which were also abundantly present amongst the highly upregulated genes in competing wild-type cells (Figure 5C). This prompted us to further investigate the exact transcriptional response induced in competing wild-type small intestinal cells and we observed enrichment of the previously reported fetal like signature (Figures 5D and S2E). Although this fetal-like response is activated in repairing intestinal epithelia, we did not find an enrichment of a Repair signature (Yui et al., 2018) in our differentially expressed genes (Figures S2F and S2G), indicating that the response of wild-type cells upon competition was different from that of an epithelium that is recovering from DSS-induced colitis. Next, we validated the observed activation of the fetal-like response by immune-fluorescence staining of SCA1. A heterogenous expression of SCA1 was detected in cancer cells, which was unaltered in mixed compared to pure cancer (Figures 5E and 5F), thereby reflecting the results of the transcriptional analysis. On the other hand, wild-type cells showed a homogenous low expression of SCA1 in unchallenged conditions, which was dramatically increased in competing cells (Figures 5E and 5G). Thus, together these data show that wild-type small intestine cells revert to a fetal-like state when challenged by competing cancer cells.

**Figure 5.**
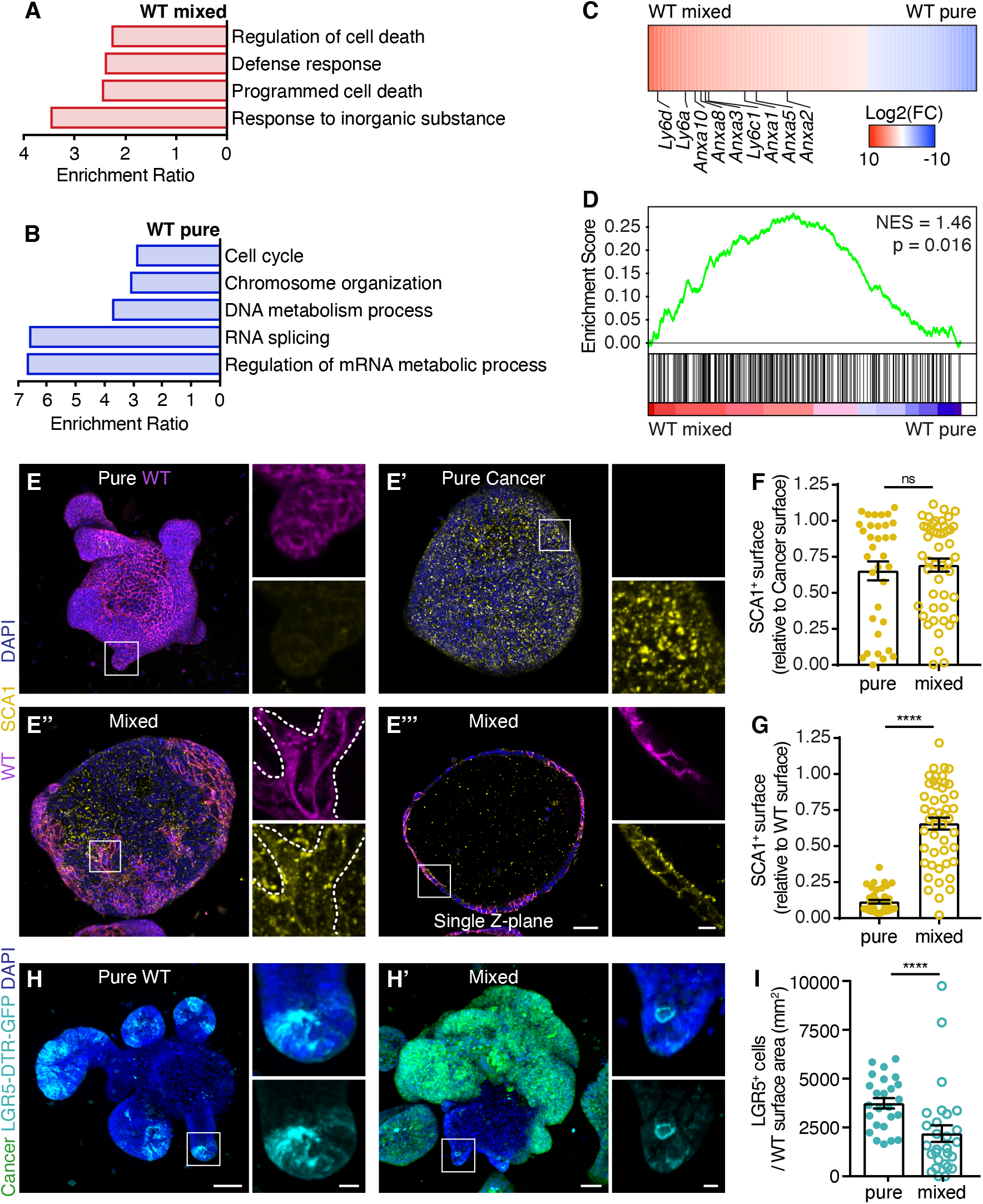
Cell competition induces a fetal-like state in wild-type cells. A, B) Gene Ontology analysis of differentially expressed genes (p<0.05) in wild-type populations that are enriched in mixed (A) and pure (B) cells. C) Heatmap of the fold change of genes that are differentially expressed in wild-type cells upon mixing (Log2). Genes of the *Ly6* and *Anxa* families are indicated. D) Gene Set Enrichment Analysis showing enrichment of a fetal signature (Yui et al., 2018) in mixed wild-type cells E-G) Representative 3D-reconstructed confocal images of pure WT (E), pure cancer (E’), mixed (E’’) organoids, and a single Z-plane of E’ (E’’’) and quantification of the SCA1+ surface relative to the total cancer (F) or wild-type (G) surface area. The organoids were stained for SCA1 (yellow), nuclei are visualized with DAPI (blue). The insets display a 3.5x magnification of the area in the white box. Each dot in (F) and (G) represents one organoid (Mean ±SEM, Non-parametric, ANOVA, multiple comparisons: p> 0.9999, n=34 & 48 organoids (F); p<0.0001, n=39 & 48 organoids (G)). H-I) Representative 3D-reconstructed confocal images of pure WT (H) and mixed (H’) organoids and quantification of the number of LGR5+ cells relative to the wild-type surface area (I). LGR5+ Intestinal stem cells (cyan) and nuclei (blue) are visualized. The insets display a 3.5x magnification of the area in the white box. Each dot in (I) represents one organoid (Mean ±SEM, Non-parametric, two-tailed, Mann-Withney: p<0.0001, two-tailed, n=25 & 28 organoids). Scale bars = 50μm, excluding magnifications in (E and H) where scale bar = 10μm

#### The cell competition-induced fetal-like state induces a loss of LGR5+ stem cells

The induction of a fetal-like state in adult intestinal epithelia has been reported to coincide with loss of intestinal stem cell (ISC) markers and removal of their niche (Nusse et al., 2018). We therefore next questioned how intestinal stem cells are affected by cancer-driven cell competition. ISCs are marked by leucine-rich-repeat-containing G-protein-coupled receptor 5 (LGR5) (Barker et al., 2007). We next derived organoids from Lgr5^DTR^ transgenic mice (Tian et al., 2011), in which the first coding exon of *Lgr5* was replaced with enhanced green fluorescent protein (eGFP) and human diphtheria toxin receptor (DTR). With the use of these Lgr5-DTR-eGFP organoids we could detect ISCs localized in crypt regions of pure wild-type organoids (Figure 5H). Upon challenging these cells with competing cancer cells, we observed a marked decrease in the number of LGR5 positive cells (Figures 5H and 5I). Thus, cancer-driven cell competition induces a cell fate transition in the surrounding wild-type epithelium that is characterized by loss of LGR5+ stem cells and adoption of a fetal-like state.

### Increased stemness prevents cell competition

So far, we have shown that wild-type intestine cells undergo a cell fate transition when exposed to competing cancer. We next wondered whether reversal of this process could disrupt cell competition. Therefore, we sought a way to interfere with the cell fate of wild-type organoids and turned to the previously described treatment with CHIR99021 and valproic acid (CV) (Yin et al., 2014). This combined inhibition of glycogen synthase kinase 3β (GSK3β) and histone deacetylases (HDAC), reported to increase self-renewal of ISCs, indeed resulted in enrichment of LGR5-positive cells in cultures after three days of treatment (Figures 6A and 6B). This coincided with loss of expression of SCA1 in mixed wild-type cells (Figures 6C and 6D), thus CV treatment can prevent the cell fate change induced by cell competition. Furthermore, time-lapse microscopy showed that inhibition of cell fate change prevented loss of wild-type cells from mixed organoids (Figures 6E and 6F). It is important to note that CV did not impact fitness of cancer cells directly, because pure cultures were not affected by the treatment (Figures S3A and S3B). Together, this shows that cell fate transition is required for elimination of wild-type cells. Interestingly, we the increased competitive potential of wild-type cells after CV treatment prevented the over proliferation of cancer cells (Figures 6G and 6H). Since the mitotic index of pure cancer cultures was not decreased by CV treatment (Figure S3C), this was not a consequence of an autonomous effect of CV treatment on cancer cells. Thus, combined these data imply that increasing stemness of wild-type cells is sufficient to prevent their elimination and neutralize the benefit that cancer obtain from competitive cell-interactions.

**Figure 6.**
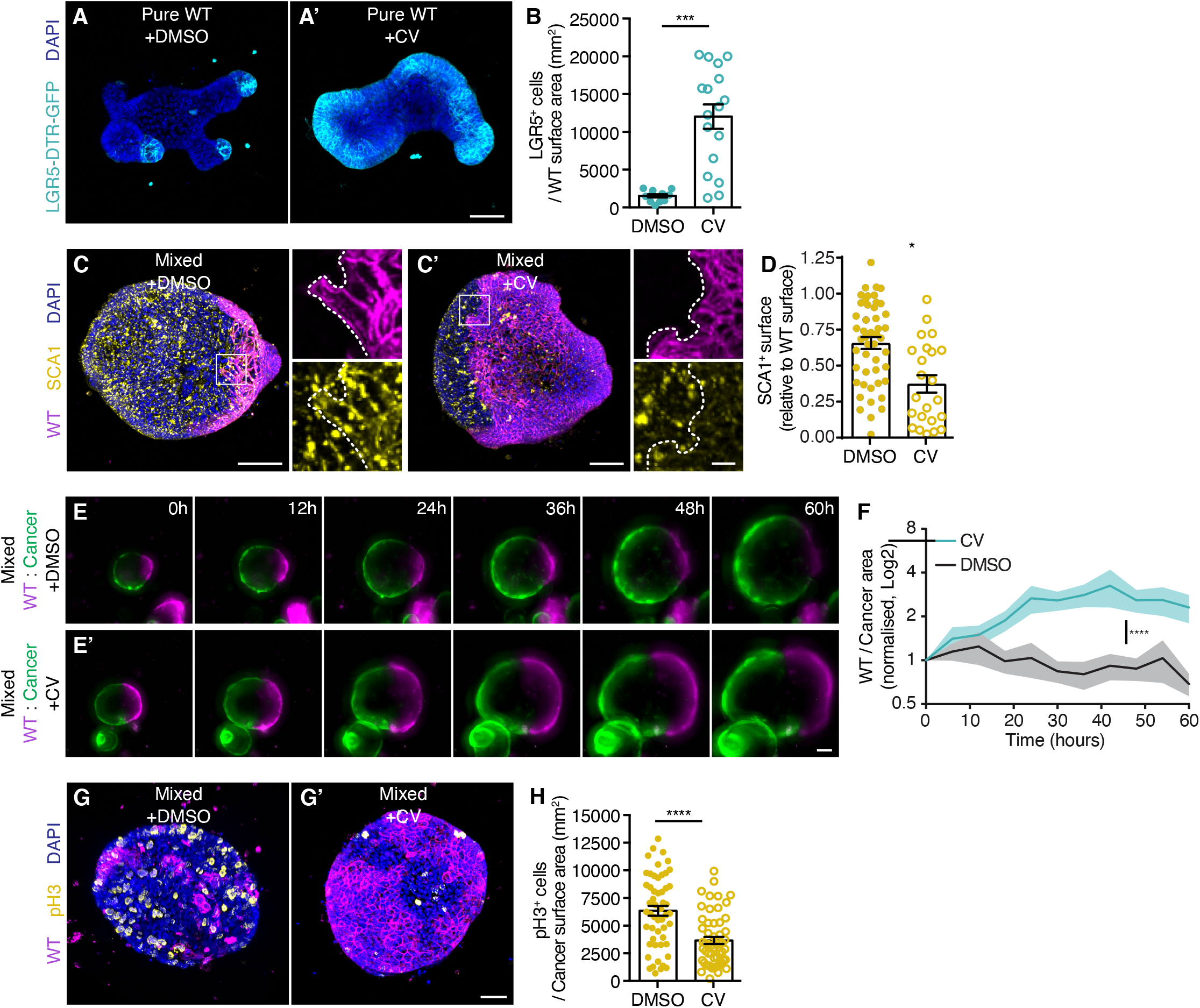
Increased stemness prevents cell competition. A) Representative 3D-reconstructed confocal images of control (A) and CV treated (A’) pure WT organoids. LGR5+ Intestinal stem cells (cyan) and nuclei (blue) are visualized. (B) displays the number of LGR5+ cells relative to the wild-type surface area, each dot represents one organoid (Mean ±SEM, unpaired T-test, two-tailed, p<0.0001, n=12 & 17 organoids). C-D) Representative 3D-reconstructed confocal images of control (C) and CV (C’) treated mixed organoids and quantification of the SCA1+ surface relative to the total wild-type surface area. The organoids were stained for SCA1 (yellow), nuclei are visualized with DAPI (blue). The insets display a 3.5x magnification of the area in the white box. Each dot in (D) represents one organoid (Mean ±SEM, Non-parametric, ANOVA, multiple comparisons: p= 0.0147, n=48 & 23 organoids). Displayed control organoids are from the same dataset used in panels 5E and 5G. E-F) Representative images of control (E) and CV (E’) treated mixed organoids by live-imaging and quantification (F) of the ratio of the area of wild-type / cancer cells in mixed organoids normalized to the start of the time-lapse on a Log2 scale (Mean ±SEM, 2-way Anova Multiple Comparisons, two-tailed, p=0.0001, n= 24 &24). Displayed control organoids are from the same dataset used in panels 4F and 4G. G-H) Representative 3D-reconstructed confocal images of control (E) and CV treated (E’) mixed organoids. Mitotic cells are marked by pH3 (yellow) and nuclei with DAPI (blue). (F) Quantification of the number of pH3+ cells relative to the cancer surface area, each dot represents one organoid (Mean ±SEM, unpaired T-test, two-tailed, p=0.0004 n=53 & 55 organoids). Scale bars = 50μm, excluding magnifications in (E) where scale bar = 10μm

### JNK signaling drives cell competition

Next, we questioned which signaling pathways could control elimination and cell fate change of wild-type cells. Therefore, we performed a transcription factor target analysis on genes that were differentially expressed in bulk mRNA sequencing. This showed that AP-1 target sites were significantly enriched in genes that were higher expressed in competing wild-type cells (Figure 7A). AP-1 transcription factors are heterodimeric proteins that are activated upon exposure to numerous stressors, such as cytokines and hypoxia (Karin and Gallagher, 2005). To further characterize activation of AP-1 we analyzed phosphorylation of cJUN, an AP-1 family member (Minden et al., 1994). We observed increased nuclear signal of cJUN-pS73 in competing wild-type cells (Figure 7B). This phosphorylation site is a substrate of the Jun N-terminal kinase (JNK), which plays a critical role in controlling both cell proliferation and death in many tissues (Karin and Gallagher, 2005; Minden et al., 1994). In addition, JNK signaling has been found to be a key regulator in many forms of cell competition (Tamori and Deng, 2011), including tumor-induced cell competition in the *Drosophila* adult intestine (Suijkerbuijk et al., 2016). We observed that treatment of mixed cultures with the selective JNK inhibitor JNK-IN-8 (Zhang et al., 2012) significantly reduced SCA1 levels in wild-type cells (Figures 7C and 7D). Suggesting that active JNK signalling is required for the cell fate state transition that is enforced by cell competition. Importantly, we found that inhibition of JNK prevents elimination of wild-type cells (Figures 7E and 7F). Thus, JNK is activated in competing cells and is required for eradication and cell fate transition of wild-type small intestine cells.

**Figure 7.**
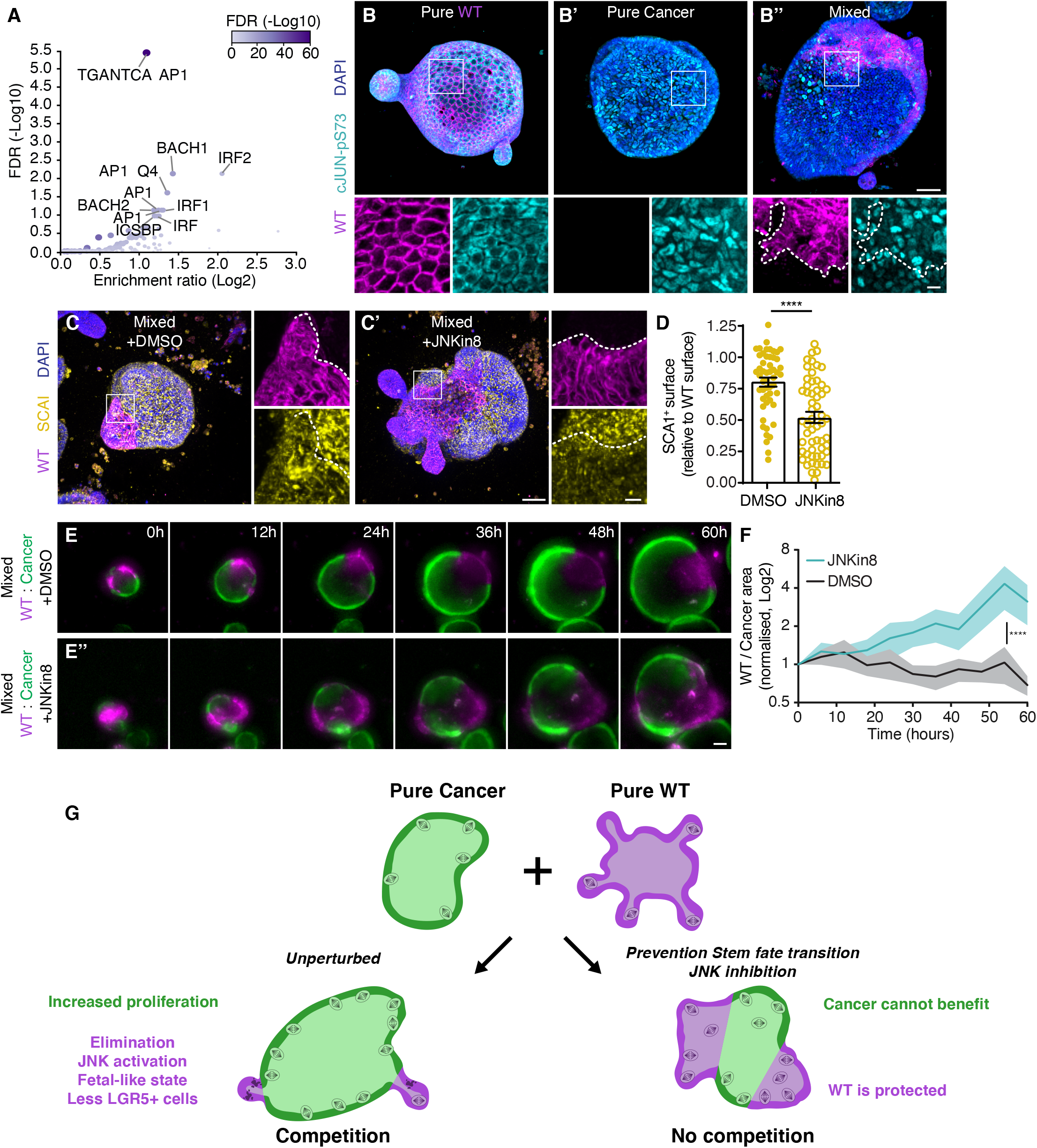
JNK signaling drives cell competition in intestinal organoids. A) Transcription factor target analysis of differentially expressed genes (p<0.05) that are enriched in mixed wild-type cells. The graph displays enrichment (Log2) and FDR (−Log10) of gene sets, significantly enriched gene sets are indicated). B) Representative 3D-reconstructed confocal images of pure WT (B), pure cancer (B’), mixed (B’’) organoids. The organoids were stained for activated cJUN (cyan), nuclei are visualized with DAPI (blue). The insets display a 3x magnification of the area in the white box. C-D) Representative 3D-reconstructed confocal images of control (C) and JNKin8 (C’) treated mixed organoids and quantification of the SCA1+ surface relative to the total wild-type surface area. The organoids were stained for SCA1 (yellow), nuclei are visualized with DAPI (blue). The insets display a 3.5x magnification of the area in the white box. Each dot in (D) represents one organoid (Mean ±SEM, Non-parametric, ANOVA, multiple comparisons: p< 0.0001, n=52 & 56 organoids). E-F) Representative images of control (E) and JNKin8 (E’) treated mixed organoids by live-imaging and quantification (F) of the ratio of the area of wild-type / cancer cells in mixed organoids normalized to the start of the time-lapse on a Log2 scale (Mean ±SEM, 2-way Anova Multiple Comparisons, two-tailed, p=0.0001, n= 24 &24). Displayed control organoids are from the same dataset used in panels 4F and 4G. G) Schematic model depicting cell competition driven growth of intestinal tumor cells in murine organoids. Cancer cells induce active elimination and cell fate transition of wild-type cells, while fuelling their oncogenic growth. Scale bars = 50μm, excluding magnifications in (B & E) where scale bar = 10μm

## Discussion

Effects of competition on cell fate have been shown in many tissues. Classical examples are neutral drift in the intestinal stem cells niche (Lopez-Garcia et al., 2010; Snippert et al., 2010) and maintenance of stem cells in the *Drosophila* testis (Sheng et al., 2009). A dual effect of cell competition on weaker cell populations, through active elimination by cell death combined with reduced stem cell renewal, has been observed under homeostasis. Both in the adult *Drosophila* midgut and in developing mouse skin, weaker cells, induced by ribosome impairment or reduced expression of *mycn*, are removed from the tissue by stronger cells through apoptosis and forced differentiation (Ellis et al., 2019; Kolahgar et al., 2015). Here, we provide the first example of a combined mechanism of forced elimination and cell fate transition in relation to cancer (Figure 8G).

Many studies have reported that tumor growth is highly context dependent. In particular, a strong correlation exists between inflammation and intestinal cancer. For example, patients with Crohn’s disease have 20-30 times higher risk of developing adenocarcinomas in the small intestine and inflammatory bowel disease is a strong risk factor for colorectal cancer (Beaugerie and Itzkowitz, 2015). Similarly, in mouse models of intestinal cancer, formation of colonic polyps is strongly enhanced by inflammation induced by infection with enterotoxigenic *Bacteroides fragilis* or treatment with dextran sodium sulfate (Tanaka et al., 2006; Wu et al., 2009). Interestingly, these are conditions in which a fetal-like response is activated in the intestine (Gregorieff et al., 2015; Nusse et al., 2018; Yui et al., 2018), similar to the here reported primitive state induced upon cancer-driven cell competition. Future efforts should be directed towards increasing understanding of this connection of a fetal-like state and tumorigenesis.

Under normal circumstances, the fetal-like response promotes regeneration of the intestinal tissue. A recent study describing the kinetics of regeneration after removal of a damaging insult has shown that reformation of a homeostatic intestinal epithelium takes approximately three weeks (Wang et al., 2019). Therefore, during chronic exposure of the epithelium to an insult, such as close proximity of a tumor, healthy tissue will never be allowed to fully recover. Interestingly, induction of a fetal-like state upon injury is not restricted to the intestinal epithelium and is also observed in multiple other tissues (Fernandez Vallone et al., 2016; Gadye et al., 2017; Lin et al., 2017). This suggests that our observation that tumors can push surrounding wild-type tissue in a primitive state can be a universal mechanism.

Cell competition can be tumor-suppressive, for example, cells expressing oncogenic H-Ras are eliminated from intestinal and pancreatic epithelia through apical extrusion (Kon et al., 2017; Sasaki et al., 2018). Furthermore, wild-type cells actively eliminate mutant aberrant foci in the skin (Brown et al., 2017). However, this effect is not solely determined by autonomous properties of the tumor but is highly context dependent. For instance, obesity induced by a high fat diet prevents cell competition-driven elimination of oncogenic cells (Sasaki et al., 2018). This illustrates how the surrounding environment dictates behavior of tumors and that tumor fitness, and thus its oncogenic potential, can be changed by external stimuli. Here, we report that JNK signaling is a major regulator of wild-type cell elimination and cell fate change, and thus overall fitness of competing healthy cells. This opens up new options of treatment, where promoting fitness of the host tissue, through JNK inhibition, can help to tip the balance towards tumor-suppressive cell competition.

## Methods

### Organoid cultures

Small intestine cancer organoids, derived from the small intestine of Villin-Cre^ERT2^*Apc*^fl/fl^*Kras*^G12D/WT^*Tr53*^fl/R172H^ mice were previously reported (Fumagalli et al., 2017b). Wild-type small intestine organoids were derived as previously described (Sato et al., 2009), from Rosa26-Cre^ERT2^::mT/mG (Muzumdar et al., 2007), Lgr5^DTR^ transgenic mice (provided by the Genentech MTA program (Tian et al., 2011)) and eCadherin-mCFP (Snippert et al., 2010) mice. All lines were cultured in drops of Cultrex PathClear Reduced Growth Factor Basement Membrane Extract Type 2 (Amsbio, 3533-005-02) in murine small intestinal organoids medium containing advanced DMEM/F12 medium (adDMEM/F12; Thermo Fisher Scientific, cat. no. 12634-010), GlutaMAX 1% (Thermo Fisher Scientific, cat. no. 35050-068), HEPES 10mM (Thermo Fisher Scientific, cat. no. 15630-056), 1x Penicillin/streptomycin (10,000 U/ml; Thermo Fisher Scientific, cat. no. 15140-122). B27 2% (Thermo Fisher Scientific, cat. no. 17504-044), N-acetylcysteine 1.25 mM (Sigma-Aldrich, cat. no. A9165), mEGF 50ng/ml (Peprotech, cat. no. 315-09), Noggin and R-spondin1 both 10% (conditioned medium prepared in house).

Enteroid monolayers were prepared as described previously (Thorne et al., 2018), in short, single cell suspensions were generated from organoids by mechanical disruption and a digest with TrypLE Express (Thermo Fisher Scientific Cat# 12605-010). Approximately 4000 WT and/or 1000 cancer cells were seeded per well of BME2 coated (0.8mg/ml) 96-well plate in medium supplemented with CHIR-99021 and Y-27632. After 24hrs, cells were washed once and cultured in murine small intestinal organoid medium for the remainder of the experiment.

3D mixed organoid cultures were prepared as follows; suspensions of small clumps of cells were generated from organoids by mechanical disruption and divided over Eppendorf vials in a 2:1 ratio (WT:cancer). Cells were concentrated by mild centrifugation, the pellet as resuspended in a small volume of murine small intestinal organoids medium and incubated at 37C for 30 minutes. Cell aggregates were plated in BME2 and cultured in murine small intestinal organoids medium.

For imaging purposes cells were plated in glass-bottom 96 well SensoPlate (Greiner Bio-One Cat#655892) or μ-Slide 8 Well chambered slides (IBIDI, cat#80827). Small molecule inhibitors were used in the following concentrations: Z-VAD-FMK (50μM, Bachem Cat# N-1510.0005), CHIR-99021 (3μM, Tocris Cat#4423), Valproic acid (1mM, Sigma Cat# PHR1061-1G), Y-2763 (10μM, Abmole Cat# M1817), JNK-IN-8 (1μM, Sigma Aldrich, Cat#SML1246).

### Immuno-fluorescence

Enteroid monolayers and organoids were fixed in 4% paraformaldehyde in PBS for 20 minutes followed by a block while permeabilizing in 5% BSA/ 0.2% Triton X100/ PBS for 30 minutes at room temperature. The stainings were performed overnight at 4°C with the following primary antibodies: anti-Cleaved Caspase-3 (Asp175) (Cell Signalling, #9661), anti-phospho-Histone H3 (Ser10) (Merck-Millipore, #06-570), anti-GFP (Abcam, #ab6673), anti-Phospho-c-Jun (Ser73) (D47G9) (Cell Signalling, #3270) and Ly-6A/E (Sca-1) (Biolegend, cat#108101). Appropriate Alexa Fluor labelled secondary antibodies (ThermoFischer Scientific) were combined with DAPI and/or Phalloidin Alexa Fluor 647 (ThermoFischer Scientific, cat# A22287). For labelling of cells in S-phase, a pulse of 10μm EdU (5-ethynyl-2'-deoxyuridine) was given one hour prior to fixation and detection was performed according to manufacturer’s guidelines before starting the immunofluorescence staining using Click-iT EdU Cell Proliferation Kit for Imaging Alexa Fluor 647 (ThermoFischer Scientific, cat# C10340).

### Microscopy

For fixed samples images were collected on an inverted Leica TCS SP8 confocal microscope (Mannheim, Germany) in 12 bit with 25X water immersion objective (HC FLUOTAR L N.A. 0.95 W VISIR 0.17 FWD 2.4 mm). Imaris software (version 9.3.1, Oxford Instruments) was used for quantification and 3D reconstructions. Images were converted to RGB using Fiji and when necessary contrasted linearly.

Sequential imaging enteriod monolayers was done using the navigator function in LasX software (Leica) on an inverted Leica TCS SP8 confocal microscope (Mannheim, Germany) in 12-bit with 25X water immersion (HC FLUOTAR L N.A. 0.95 W VISIR 0.17 FWD 2.4 mm). Merged images were rotated, aligned and cropped in Photoshop (Adobe) and when necessary contrasted linearly.

Time-lapse microscopy of enteroid monolayers was performed on a Leica TCS SP5 confocal microscope (Mannheim, Germany) in 12-bit with 20X dry immersion objective (HCX PL APO CS 20.0×0.70 DRY UV). Images were converted to RGB, cropped and when necessary contrasted linearly in FIJI.

Time-lapse microscopy of 3D organoids was performed on an AxioObserver widefield microscope (Zeiss) equipped with an Orca FLASH 4.0 V3 grayscale sCMOS-camera (Hamamatsu) and using a 10X dry objective (N.A. 0.30 EC Plan-Neofluar Ph1). Whole drops of organoids were followed up to 90 hours and images were collected in 16-bit with a 6-hour time interval. ZEN software (Zeiss) was used to stitch mosaic images. Quantification and image processing were performed using FIJI; images were converted to RGB, cropped and when necessary contrasted linearly.

### Statistical analysis

Statistics were performed using GraphPad Prism. Paired or unpaired t-test was used when data showed normal distribution (verified with normality tests, provided by GraphPad Prism), whereas Mann-Whitney U test was used for data that did not display parametric distribution. Adoption of one statistical test or the other is indicated for each experiment in the Figure legend.

### Transcriptomics

For gene expression analysis, pure and mixed cells were cultured in 6-well plates and FACS sorted 3 days after mixing on a FACS Jazz system (BD). After a FSC/SSC gate doublets were excluded, live cells (DAPI negative) were sorted based on mTomato (wild-type cells) and Dendra2 (cancer cells) expression. Purity was determined by microscopy and data were analyzed using FlowJo.

Total RNA was extracted using the standard TRIzol (Invitrogen) protocol and used for library preparation and sequencing. mRNA was processed as described previously, following an adapted version of the single-cell mRNA seq protocol of CEL-Seq (Hashimshony et al., 2012; Simmini et al., 2014). In brief, samples were barcoded with CEL-seq primers during a reverse transcription protocol and pooled after second strand synthesis. The resulting cDNA was amplified with an overnight *In vitro* transcription reaction. From this amplified RNA, sequencing libraries were prepared with Illumina Truseq small RNA primers. Paired-end sequencing was performed on the Illumina Nextseq500 platform. Read 1 was used to identify the Illumina library index and CEL-Seq sample barcode. After quality control and adaptor removal, read 2 was aligned to the hg19 human RefSeq transcriptome using BWA (Li and Durbin, 2010). Reads that mapped equally well to multiple locations were discarded. Reads were quantified with featureCounts to generate read counts for each gene based on the gene annotation from Ensemble. Differential gene expression was analyzed, based on featureCounts results, using DEseq2 version 1.22.2 (Love et al., 2014).

Gene Ontology analysis was performed with Webgestalt (Liao et al., 2019) (http://www.webgestalt.org/), with the parameters; Method: Over-representation Analysis (ORA); Organism: mmusculus; Enrichment Categories: geneontology_Biological_Process; FDR Method: BH; Significance Level: Top 10; Redundancy reduction correction using Weighted set cover algorithm, FDR ≤ 0.05.

The heat map was generated using Morpheus developed by the Broad Institute (https://software.broadinstitute.org/morpheus) and scaled in Adobe Illustrator.

Gene Set Enrichment Analysis was performed with GSEA software developed by UC San Diego and Broad Institute (Mootha et al., 2003; Subramanian et al., 2005) (https://www.gsea-msigdb.org/gsea/index.jsp). Standard parameters were used with a pre-ranked dataset of differentially expressed genes in wild-type cells (p<0.05). Gene Sets that were used for comparison were the previously published ‘mFetal’ and ‘mRepair’ (Yui et al., 2018).

Transcription Factor target analysis was performed with Webgestalt (Liao et al., 2019) (http://www.webgestalt.org/), with the parameters; Method: Over-representation Analysis (ORA); Organism: mmusculus; Enrichment Categories: network_Transcription_Factor_target; FDR Method: BH; Significance Level: Top 10.

## Acknowledgements

We thank Marjolijn Mertz, Lenny Brocks and the NKI BioImaging facilty and Anko de Graaff and the Hubrecht Imaging Centre for technical support with imaging. Evelyne Beerling and Tim Schelfhorst for technical support. Judith Vivié for help with mRNA sequencing. Kim Jensen for sharing gene enrichment datasets, Frederic J. de Sauvage for sharing the Lgr5-DTR-GFP mouse and members of the van Rheenen group for critically reading the manuscript.

This work was financially supported by Dutch Cancer Society Fellowship BUIT-2013-5847 (to S.J.E.S.) and Young Investigator Grant 11491 (to S.J.E.S. and A.K.G) and the CancerGenomics.nl (to J.v.R.), European Research Council Grant CANCER-RECURRENCE 648804 (to J.v.R.), the Doctor Josef Steiner Foundation (to J.v.R) and the European Union's Horizon 2020 research and innovation program under the Marie Sklodowska-Curie grant agreement No 642866 (to J.v.R). This work was also supported by Cancer Research UK core grant to the Beatson Institute core (A17196), CRUK core funding A21139 and CRUK ACRCelerate A26825 (both to O.J.S.). This manuscript was edited by Life Science Editors.

## Author contributions

S.J.E.S and A.K.G. performed the experiments. H.L.Q. performed mRNA sequencing analysis, A.F., O.J.S and J.v.R. contributed knowledge and reagents. S.J.E.S. designed the experiments, supervised the study and wrote the manuscript together with J.v.R., and the manuscript was approved by all authors. Correspondence and requests for materials should be addressed to S.J.E.S.

## Competing Interests statement

The authors declare no competing interests.

## Notes

### Competing Interest Statement

The authors have declared no competing interest.

