## Supplementary Figures for "Active elimination of intestinal cells drives oncogenic growth in organoids"

#### Supplementary Information

### Cell competition drives growth of cancer cells through forced cell death and cell fate change in intestinal organoids

*Krotenberg Garcia et al*

### Krotenberg Garcia et al, Supplementary Figure 1

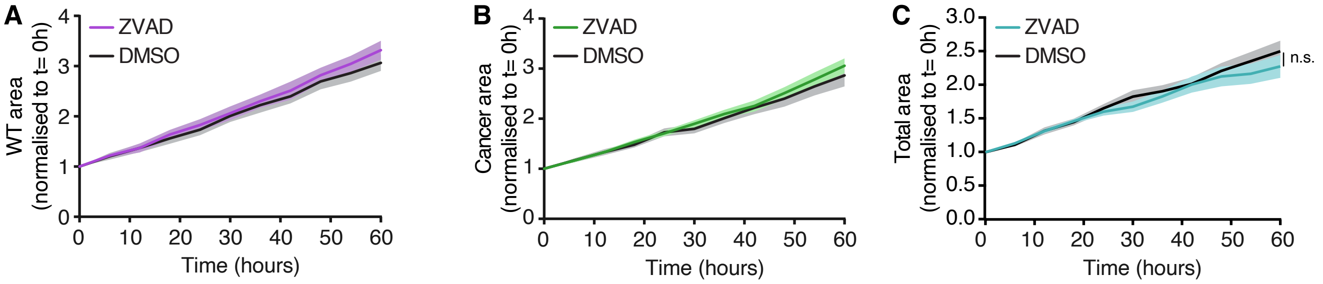


**Supplementary Figure 1 – Elimination of wild-type cells is driven by apoptosis**

A-C) Analysis of growth of pure WT (A), pure cancer (B) and mixed (C) organoids treated with DMSO control or Z-VAD-FMK by live-imaging. Graph displays the total covered area within organoids normalised to the start of the time-lapse (Mean ±SEM, 2-way ANOVA Multiple Comparisons, two-tailed, p=0.2383 n= 12 & 12 (A); p=0.4441 n= 22 & 22 (B); p=0.1455 n=30 & 30(C)).

### Krotenberg Garcia et al, Supplementary Figure 2

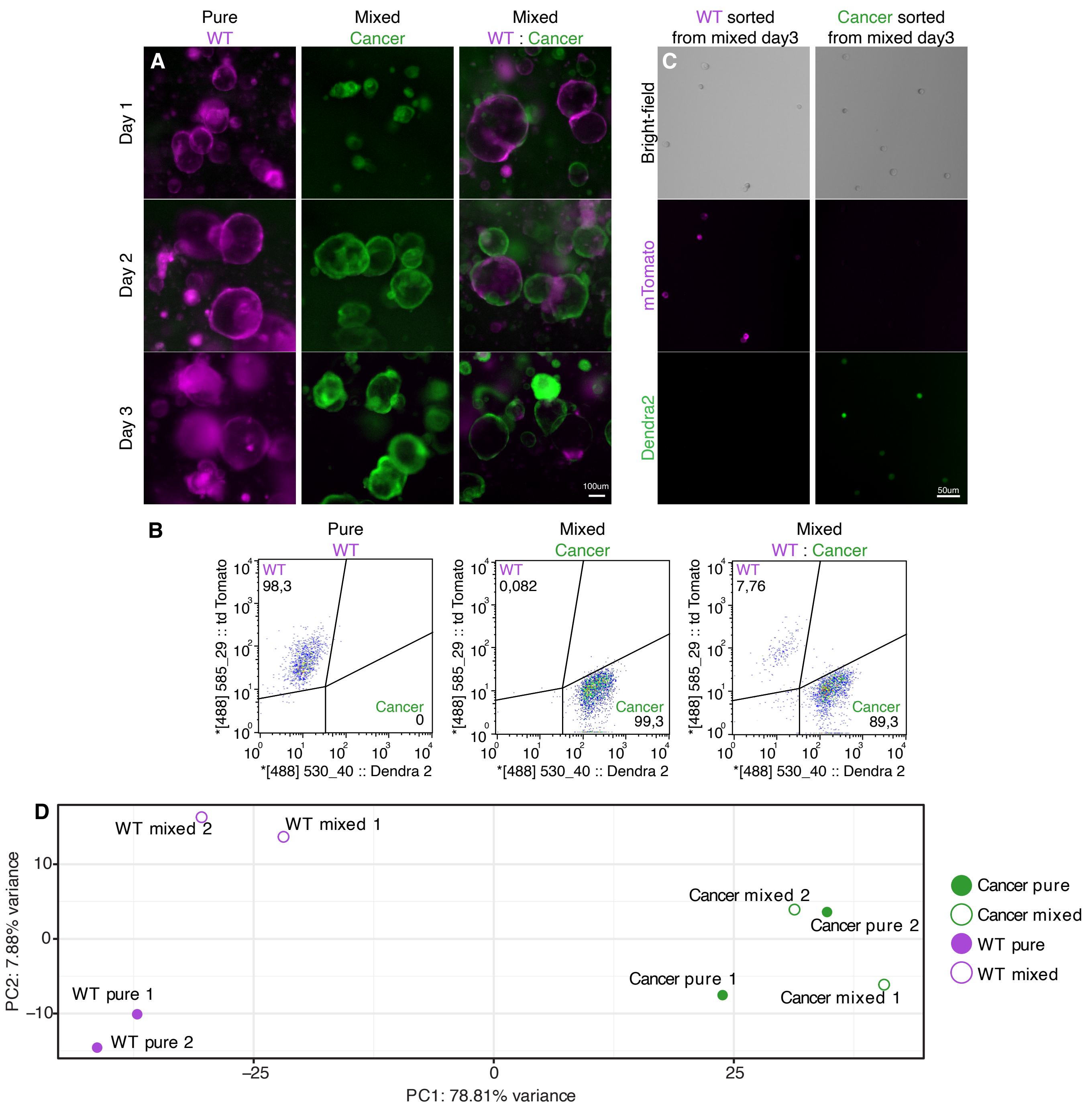


**E**

| GeneSet | mFetal |
|---|---|
| Enrichment Score (ES) | 0.27931073 |
| Normalized Enrichment Score (NES) | 1.4551047 |
| Nominal p-value | 0.016477859 |
| FDR q-value | 0.056112226 |
| FWER p-Value | 0.052 |

**F**

| GeneSet | mRepair |
|---|---|
| Enrichment Score (ES) | 0.24593316 |
| Normalized Enrichment Score (NES) | 1.1627909 |
| Nominal p-value | 0.2491731 |
| FDR q-value | 0.2249499 |
| FWER p-Value | 0.374 |

**G**

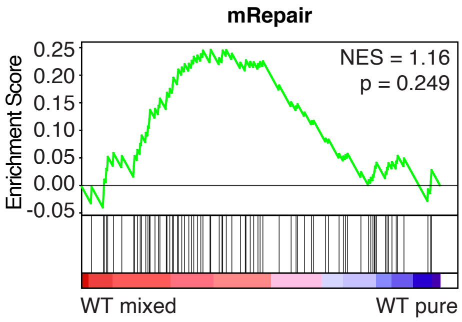


*(legends on next page)*

**Supplementary Figure 2 - Cell competition induces a fetal-like state in WT cells**

A-C) Flow cytometry sorting of wild-type and cancer cells from pure and mixed cultures. A) Representative images of pure and mixed cultures 1, 2 and 3 days after plating, scale bar = 50µm. B-C) Analysis of cells after sorting, graphs in B show an analysis of 10.000 cells, numbers in the corners display the percentage of sorted cells. Representative images of sorted cells are shown in (C) Scale bars = 100µm.

D-G) Gene expression analysis of wild-type and cancer cells in pure and mixed conditions. D) displays a principal component analysis of all sample. D) Parameters of a gene Set Enrichment Analysis showing enrichment of a fetal signature 18 in mixed wild-type cells. F-G) Parameters and graph of a gene Set Enrichment Analysis showing no significant enrichment of a repair signature 18 in mixed wild-type cells.

### Krotenberg Garcia et al,  Supplementary Figure 3

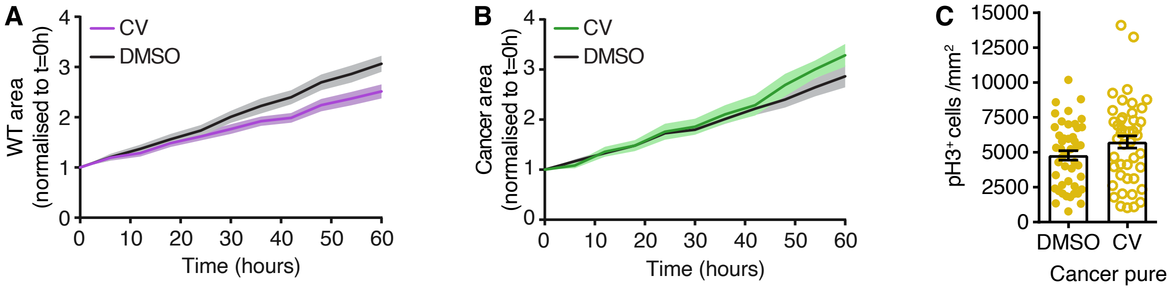


**Supplementary Figure 2 - Increased stemness prevents cell competition**

A-B) Analysis of growth of pure WT (A) and cancer (B) organoids treated with DMSO control or CV by live-imaging. Graph displays the covered area within organoids normalised to the start of the time-lapse (Mean ±SEM, 2-way Anova Multiple Comparisons, two-tailed, p=0.0001 n= 12 &12(A); p=0.1268 n= 22 &12(B).

C) Graph displays the number of pH3+ cells relative to the cancer surface area, each dot represents one organoid (Mean ±SEM, unpaired T-test, two-tailed, p=0.0864, n=46 & 46 organoids).

### Supplementary Movies

**Movie 1:**
Time-lapse series of a competing enteroid monolayer, arrow heads indicate examples of wild-type cells that are shrinking and being eliminated.

**Movie 2:**
Time-lapse series of a competing enteroid monolayer, arrow heads indicate examples of wild-type cells that are shrinking and being eliminated.

**Movie 3:**
Time-lapse series of pure wild-type intestinal organoid.

**Movie 4:**
Time-lapse series of mixed intestinal organoid.

**Movie 5:**
3D-reconstruction of confocal images and mixed organoid. The actin cytoskeleton is stained with Phalloidin (yellow), nuclei with DAPI (blue).
